# *Plasmodium falciparum* malaria drives epigenetic reprogramming of human monocytes toward a regulatory phenotype

**DOI:** 10.1101/2020.10.21.346197

**Authors:** Rajan Guha, Anna Mathioudaki, Safiatou Doumbo, Didier Doumtabe, Jeff Skinner, Gunjan Arora, Shafiuddin Siddiqui, Shanping Li, Kassoum Kayentao, Aissata Ongoiba, Judith Zaugg, Boubacar Traore, Peter D. Crompton

## Abstract

In malaria-naïve children and adults, *Plasmodium falciparum*-infected red blood cells (*Pf*-iRBCs) trigger fever and other symptoms of systemic inflammation. However, in endemic areas where individuals experience repeated *Pf* infections over many years, the risk of *Pf*-iRBC-triggered inflammatory symptoms decreases with cumulative *Pf* exposure. The molecular mechanisms underlying these clinical observations remain unclear. Age-stratified analyses of monocytes collected from uninfected, asymptomatic Malian individuals before the malaria season revealed an inverse relationship between age and *Pf*-iRBC-inducible inflammatory cytokine (IL-1β, IL-6 and TNF) production, whereas Malian infants and malaria-naïve U.S. adults produced similarly high levels of inflammatory cytokines. Accordingly, monocytes of Malian adults produced more IL-10 and expressed higher levels of the regulatory molecules CD163, CD206, Arginase-1 and TGM2. These observations were recapitulated in an *in vitro* system of monocyte to macrophage differentiation wherein macrophages re-exposed to *Pf*-iRBCs exhibited attenuated inflammatory cytokine responses and a corresponding decrease in the epigenetic marker of active gene transcription, H3K4me3, at inflammatory cytokine gene loci. Together these data indicate that *Pf* induces epigenetic reprogramming of monocytes/macrophages toward a regulatory phenotype that attenuates inflammatory responses during subsequent *Pf* exposure. These findings also suggest that past malaria exposure could mitigate monocyte-associated immunopathology induced by other pathogens such as SARS-CoV-2.

**Author Summary:** The malaria parasite is mosquito-transmitted and causes fever and other inflammatory symptoms while circulating in the bloodstream. However, in regions of high malaria transmission the parasite is less likely to cause fever as children age and enter adulthood, even though adults commonly have malaria parasites in their blood. Monocytes are cells of the innate immune system that secrete molecules that cause fever and inflammation when encountering microorganisms like malaria. Although inflammation is critical to initiating normal immune responses, too much inflammation can harm infected individuals. In Mali, we conducted a study of a malaria-exposed population from infants to adults and found that participants’ monocytes produced less inflammation as age increases, whereas monocytes of Malian infants and U.S. adults, who had never been exposed to malaria, both produced high levels of inflammatory molecules. Accordingly, monocytes exposed to malaria in the laboratory became less inflammatory when re-exposed to malaria again later, and these monocytes ‘turned down’ their inflammatory genes. This study helps us understand how people become immune to inflammatory symptoms of malaria and may also help explain why people in malaria-endemic areas appear to be less susceptible to the harmful effects of inflammation caused by other pathogens such as SARS-CoV-2.

## Introduction

*Plasmodium falciparum* infection in non-immune individuals can result in severe, life-threatening malaria when *P*. *falciparum-infected* red blood cells (*Pf*-iRBCs) trigger systemic inflammation [1],[2] and sequester in blood vessels of vital organs [3]. Conversely, in areas of intense malaria transmission, *P*. *falciparum*-infected individuals are commonly asymptomatic [4], even when parasitemia exceeds that which predictably induces fever and other inflammatory symptoms in non-immune individuals. Non-sterilizing, clinical immunity to blood-stage malaria can be acquired with repeated infections over years and is associated with the acquisition of *P*. *falciparum*-specific humoral and cellular adaptive immune responses [5, 6]. The relatively inefficient acquisition of adaptive immunity that protects from malaria has been ascribed to the extensive genetic diversity of *P*. *falciparum parasites* [7], the clonal variation in proteins the parasite exports to the erythrocyte surface [8], and dysregulation of B and T cell responses [9, 10].

Less is known about the impact of cumulative *P*. *falciparum* exposure on cells of the innate immune system, such as monocytes and macrophages, and how this may relate to the acquisition of clinical immunity. During *Plasmodium* blood-stage infection, circulating blood monocytes and tissue macrophages perform crucial effector functions that contribute to host defense against malaria including cytokine production, phagocytosis of infected erythrocytes, and antigen presentation [11]. However, excessive production of pro-inflammatory cytokines such IL-1β, IL-6 and TNF by monocytes/macrophages can result in systemic inflammation that causes fever and other disease manifestations of malaria [11, 12].

Recent studies have shown that various immune perturbations can epigenetically and metabolically reprogram monocytes/macrophages, such that after cells return to a non-activated state, their response to subsequent challenges is altered [13]. Depending on the nature of the initial immune perturbation, the subsequent response of monocytes/macrophages may be diminished (tolerance) or enhanced (trained immunity) relative to the primary response [13]. It has been shown that immune training of monocytes can be generated at the level of myeloid progenitors in the bone marrow [14], which could explain how monocytes, which survive in circulation for only 1-7 days [15], could exhibit a ‘memory’ phenotype for 3-12 months after the primary stimulus [13].

Therefore, we hypothesized that *P*. *falciparum* exposure is associated with functional changes in circulating monocytes that persist in the absence of ongoing malaria exposure when monocytes have returned to a non-activated steady state. More specifically, given the long-standing clinical observation that individuals become ‘tolerant’ to the inflammatory effects of blood-stage malaria [16], we hypothesized that an inverse relationship exists between age (a surrogate for cumulative malaria exposure in endemic areas) and *Pf*-iRBC-inducible inflammatory cytokine production from monocytes at their non-activated steady state.

To test this hypothesis, we analyzed the phenotype and function of monocytes obtained from an age-stratified cohort study in Mali that spans infancy to adulthood. Monocytes were collected cross-sectionally from asymptomatic, uninfected study volunteers at the end of the 6-month dry season, which is a period of negligible *P*. *falciparum* transmission. In addition, we adapted an in vitro system of monocyte to macrophage differentiation to directly investigate the impact of *P*. *falciparum* blood-stage parasites on monocytes/macrophages at the molecular level.

## Results

### Malaria exposure associates with reduced production of Pf-inducible inflammatory cytokines and increased production of IL-10 from monocytes

To analyze the relationship between malaria exposure and monocyte function we isolated monocytes from subjects enrolled in an age-stratified cohort in Mali and also from healthy malaria-naïve U.S. adults as controls. A detailed description of the Kalifabougou cohort has been published [17]. Malian subjects ranged from 4-6-month-old infants born during the six-month dry season, when malaria transmission is negligible [18], to adults exposed to a lifetime of repeated *P*. *falciparum* infections. All Malian subjects in this cross-sectional analysis were negative for *P*. *falciparum* infection by PCR at the time of blood collection, which occurred just before the 6-month malaria season. To simulate re-exposure to *P*. *falciparum* blood-stage parasites in vitro, isolated monocytes were co-cultured with P. falciparum-infected red blood cells (*Pf*-iRBCs) at a ratio of 1:30 (monocytes:*Pf*-iRBCs) for 24 hours [19]. Secreted cytokines were measured in supernatants by bead-based multiplex arrays. Among the Malian subjects we observed an inverse relationship between age and the production of the inflammatory cytokines TNF, IL-1β and IL-6, whereas Malian infants and malaria-naïve U.S. adults produced similarly high levels of inflammatory cytokines (Figure 1 A-C). In contrast, production of the anti-inflammatory cytokine IL-10 increased with age among Malian subjects, whereas Malian infants and malaria-naïve U.S. adults produced similarly low levels of IL-10 (Figure 1D). Taken together, these data suggest that cumulative malaria exposure, or exposure to other factors associated with malaria transmission, skews monocytes toward a regulatory phenotype characterized by decreased *P*. *falciparum*-inducible production of inflammatory, pyrogenic cytokines and increased production of the anti-inflammatory cytokine IL-10, consistent with the epidemiological observation that febrile malaria risk decreases with cumulative malaria exposure in this cohort [17], and in areas of intense malaria transmission more generally [20].

**Figure 1.**
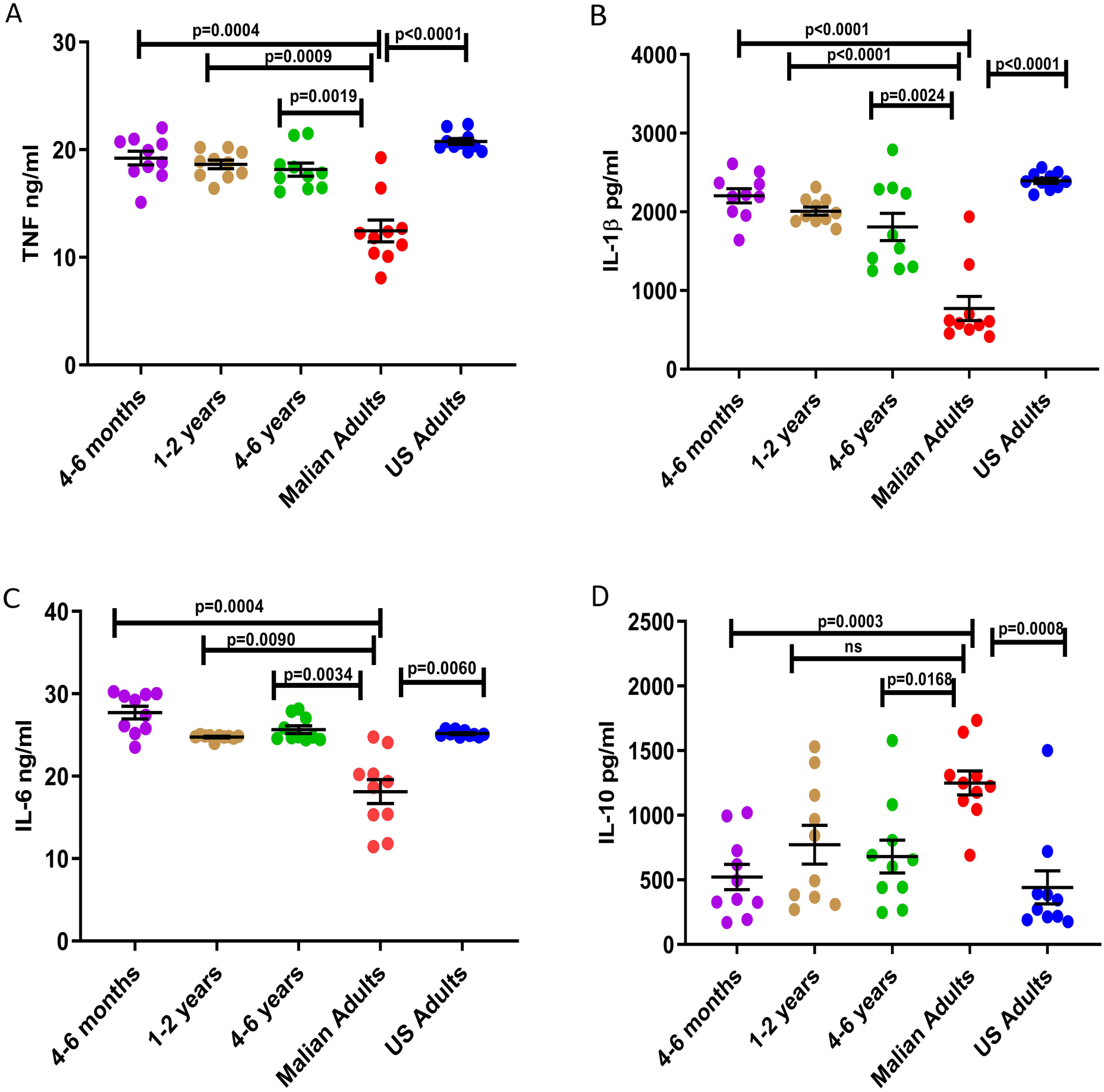
Monocytes of Malian adults exhibit reduced *P*. *falciparum*-inducible inflammatory cytokine production and increased IL-10 production. PBMCs were collected cross-sectionally from an age-stratified cohort in Mali before the malaria season when all subjects were negative for *Pf* infection by PCR; and also, from malaria-naïve U.S. adults. Monocytes were isolated from thawed PBMCs and stimulated with iRBC lysate at a ratio of 1 monocyte to 30 iRBCs. After 24 hours, cell culture supernatants were analyzed to quantify secreted (**A**) TNF, (**B**) IL-1β, (**C**) IL-6 and (**D**) IL-10. Means ± SEM are shown. Data were analyzed by the Brown Forsythe and Welch ANOVA test followed by Dunnett’s T3 multiple comparison test. Level of significance between groups are indicated by P values.

### Monocytes of malaria-exposed adults skew toward a regulatory profile phenotypically and transcriptionally

Next, we examined the molecular basis of malaria-associated skewing of monocytes toward a regulatory phenotype by comparing the phenotypic and transcriptional profiles of monocytes collected before the malaria season from Malian children and adults, as well as malaria-naïve U.S. adults. PBMCs of Malian children (aged 4-6 years; n=9) and adults (n=9), as well as U.S. adults (n=7) were analyzed *ex vivo* by flow cytometry for surface expression of the myeloid cell markers CD14, CD16, CD86, CD163, CD206 and HLA-DR [21] gated on live monocytes. Visualization of the flow cytometry data by distributed stochastic neighbor embedding (t-SNE) analysis [22] revealed that monocytes of Malian adults expressed higher levels of the regulatory or alternatively activated (M2) monocytes/macrophage markers CD163 and CD206 compared to Malian children and U.S. adults (Figure 2A); and CD163 and CD206 expression was largely confined to the CD14+ and CD14+CD16+ monocyte clusters (Figure 2B). Consistent with the t-SNE visualization, MFI values of CD163 and CD206 were significantly higher on monocytes of Malian adults compared to Malian children and U.S. adults (Figure 2C and D).

**Figure 2.**
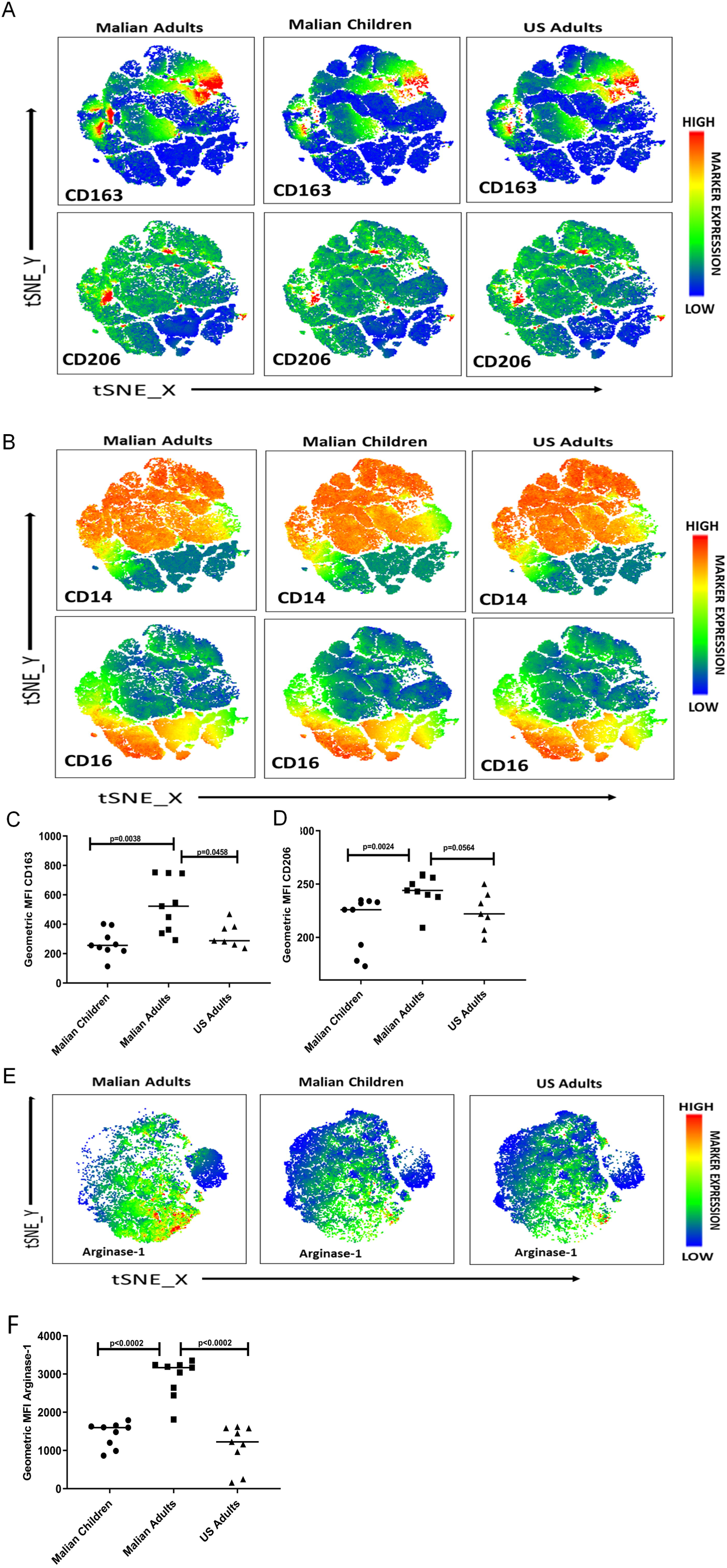
Monocytes of Malian adults upregulate markers of M2 regulatory monocytes/macrophages. (**A-D**) Monocytes from Malian children (aged 4-6 years; n=9) and adults (n=9) before the malaria season, as well as healthy malaria-naïve U.S. adults (n=7) were analyzed *ex vivo* by flow cytometry for surface expression of CD14, CD16, CD163 and CD206. (**A, B**) t-SNE analysis of monocytes for all subjects in each group. Expression of each marker is indicated by a color scale. MFI of (**C**) CD163 and (**D**) CD206 surface expression on manually gated live monocytes. (**E,F**) PBMCs were stimulated with *Pf*-iRBCs and analyzed for arginase-1 expression in monocytes intracellularly. (**E**) Representative t-SNE plots showing expression of Arginase-1 in the clusters in three different groups of monocytes. Expression of Arginase-1 is indicated by a color scale. (**F**). Expression MFI level of intracellular Arginase-1 on manually gated live monocytes comparing three different groups. Data were analyzed by the Mann-Whitney test with Bonferroni adjustment, and the level of significance between groups is indicated by P values.

M2 regulatory monocytes/macrophages are known to produce high levels of arginase 1 [23]. Therefore we further tested the hypothesis that cumulative malaria exposure is associated with skewing of monocytes towards an M2 regulatory phenotype by stimulating PBMCs of Malian children (aged 4-6 years; n=9) and adults (n=9), as well as U.S. adults (n=9) with *Pf*-iRBCs for 18 hours and quantifying intracellular arginase-1 in monocytes by flow cytometry. Visualization of the flow cytometry data by t-SNE analysis showed that monocytes of Malian adults produced higher levels of arginase 1 in response to *Pf*-iRBC stimulation compared to Malian children and U.S. adults (Figure 2E). Consistent with the t-SNE plots, MFI values of arginase 1 were significantly higher within monocytes of Malian adults compared to Malian children and U.S. adults (Figure 2F).

To further investigate the molecular basis of malaria-associated skewing of monocytes toward an M2 regulatory type, we conducted whole genome RNA-seq analysis of monocytes isolated from Malian children and adults. RNA-seq was performed on unstimulated monocytes (n=4 from each age group) as well as monocytes that had been stimulated with *Pf*-iRBCs for 24 hours (n=8 from each age group) to simulate re-exposure to P. falciparum blood-stage parasites. Monocytes were stimulated with *Pf*-iRBCs at a ratio of 1:5 (monocytes:*Pf*-iRBCs) to reduce *P*. *falciparum* nucleic acid during RNA sequencing. Principal components analysis of the RNA-seq data showed segregation of transcription profiles based on age—an effect that became more pronounced following stimulation with *Pf*-iRBCs (Figure 3A). Here we focused on differential gene expression between monocytes of children versus adults following *Pf*-iRBC stimulation. Consistent with the analysis of secreted cytokines (Figure 1), Malian children had significantly higher expression of the genes encoding TNF [log2 fold change (FC) 1.8, Benjamini-Hochberg (BH) adjusted p value = 0.0001] and IL6 (FC 2.3, BH p = 0.007) compared to Malian adults (Figure 3B and C; Supplemental Table). In addition, the expression of *TLR5, TLR7, CXCL9, CXCL10, NLRP1, NLRP3, FCGR3A, PTX3* and various HLA molecules was significantly upregulated in children relative to adults (Figures 3B-E; Supplemental Table for FC and adjusted p values). Consistent with the blunted pro-inflammatory cytokine responses that we observed from monocytes of Malian adults in response to *Pf*-iRBC stimulation (Figure 1), expression of NFKB1, a positive regulator of inflammation, was downregulated (FC −3.8, BH p = 4.6 E-26) in monocytes of adults relative to children (Figure 3C; Supplemental Table), while expression of the multifunctional enzyme transglutaminase-2 (*TGM2*), a known marker of the M2 regulatory phenotype, was significantly higher (FC 7.5, BH p = 1.7 E-79) in monocytes of adults versus children (Figures 3B and D; Supplemental Table). Accordingly, expression of the chemokines *CCL22* (FC 5.0, BH p = 9.2 E-21) and *CCL24* (FC 9.6, BH p = 3.8 E-21) were upregulated in adult monocytes, consistent with the chemokine repertoire of M2 regulatory monocytes/macrophages [24].

**Figure 3.**
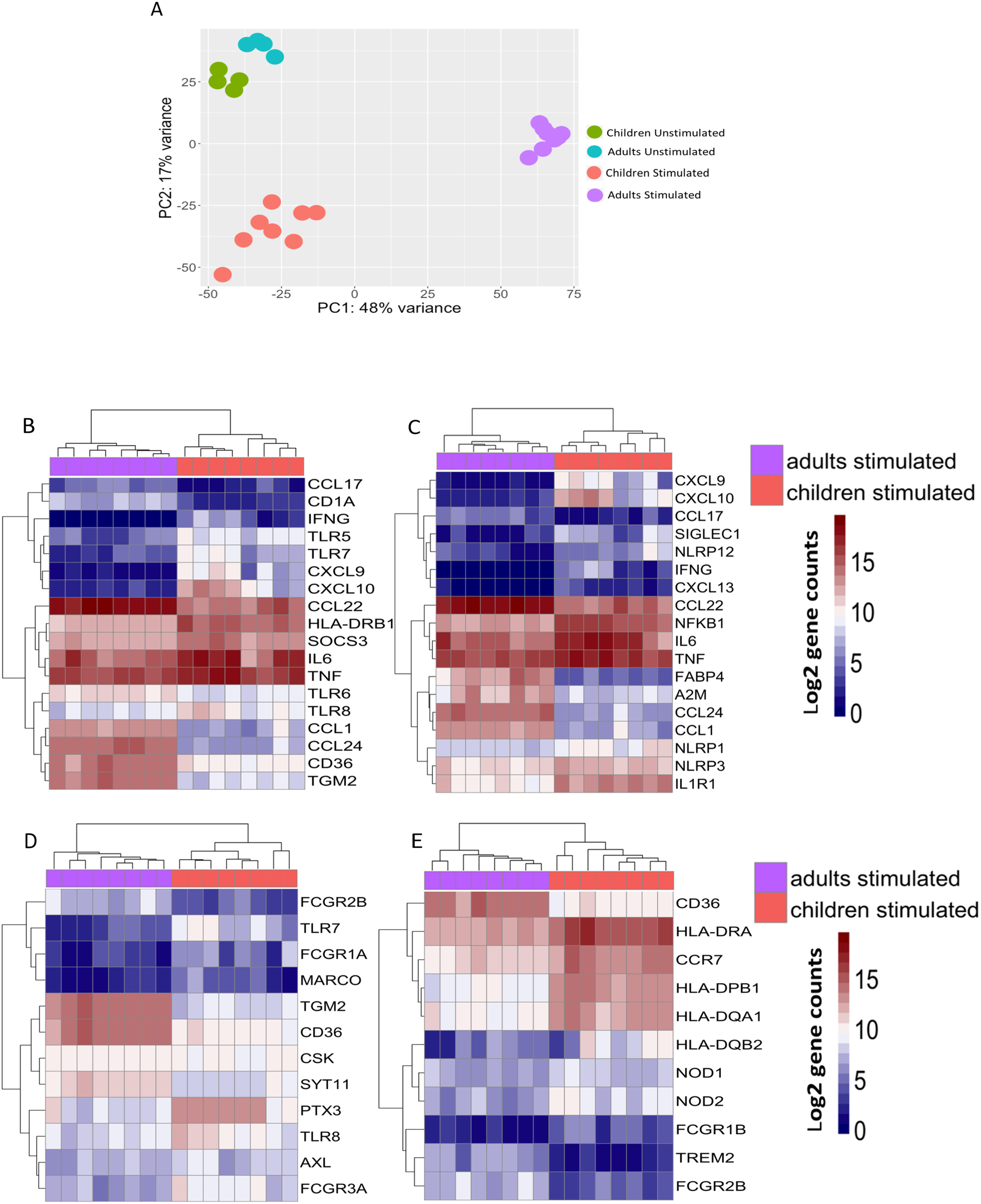
RNA-seq analysis of monocytes after *Pf*-iRBC stimulation reveals a regulatory signature in Malian adults that is distinct from children. Isolated monocytes from Malian children (n=8) and adults (n=8) were stimulated *in vitro* with *Pf*-iRBC lysate at a monocyte: *Pf*-iRBC ratio of 1:5 for 24 hours and then total RNA was isolated for sequencing. From separate subjects (n=4 subjects in each age group), total RNA from unstimulated monocytes was also sequenced. (**A**) Principal-component analysis of log2-normalized gene counts across all samples. (**B-E**) Heatmaps representing log2-normalized gene counts of *Pf*-iRBC stimulated monocytes from Malian children and adults. Each column represents one individual sample. Heatmaps represent gene sets with pre-specified functions: (**4B**) M1/M2 monocyte/macrophage signature, (**4C**) inflammation, (**4D**) phagocytosis, and (**4E**) antigen processing and presentation.

Together these data indicate that monocytes of Malian adults are skewed toward a regulatory phenotype, whereas monocytes of Malian children more closely resemble those of malaria-naïve U.S. adults, suggesting that cumulative malaria exposure, or other factors associated with malaria transmission, and not age *per se*, drives functional changes in monocytes that persist in uninfected individuals.

### Pre-exposure of monocytes to *P*. *falciparum* blunts subsequent inflammatory responses to *P*. *falciparum* and LPS

The Malian adults in this study whose monocytes skew toward a regulatory phenotype have been exposed to a lifetime of repeated *P*. *falciparum* infections, but whether *P*. *falciparum per se* can drive this phenotype remains unclear. To more directly test this hypothesis, and to further dissect the molecular mechanisms underlying our ex vivo observations, we adapted an in vitro model of monocyte to macrophage differentiation [25] to incorporate exposure to *P*. *falciparum* blood-stage parasites (Figure 4). Briefly, elutriated monocytes from healthy U.S. adults were incubated for 24 hours with medium alone, uninfected red blood cells (RBC) or *Pf*-iRBC. At 24 hours, supernatants and cells were collected from some replicate wells for cytokine analysis, while monocytes in other replicate wells were washed and incubated for 3 additional days in human serum and medium to allow monocytes to differentiate into macrophages (Mf). On day 5, the three populations of macrophages were either harvested for ChIP and cytokine gene expression analysis or re-stimulated with *Pf*-iRBCs or lipopolysaccharide for 24 hours prior to measuring cytokines in supernatants. Of note, on day 5, cell frequency and viability did not differ significantly between macrophages in medium alone (Mf) or following co-culture with RBCs (RBC-Mf) or Pf-iRBCs (*Pf*-iRBC-Mf) (Supplementary Figure 2). Also of note, titration experiments determined that a monocyte:*Pf*-iRBC ratio of 1:15 optimally balanced cytokine production and monocyte viability in this model (Supplementary Figure 1).

**Figure 4.**
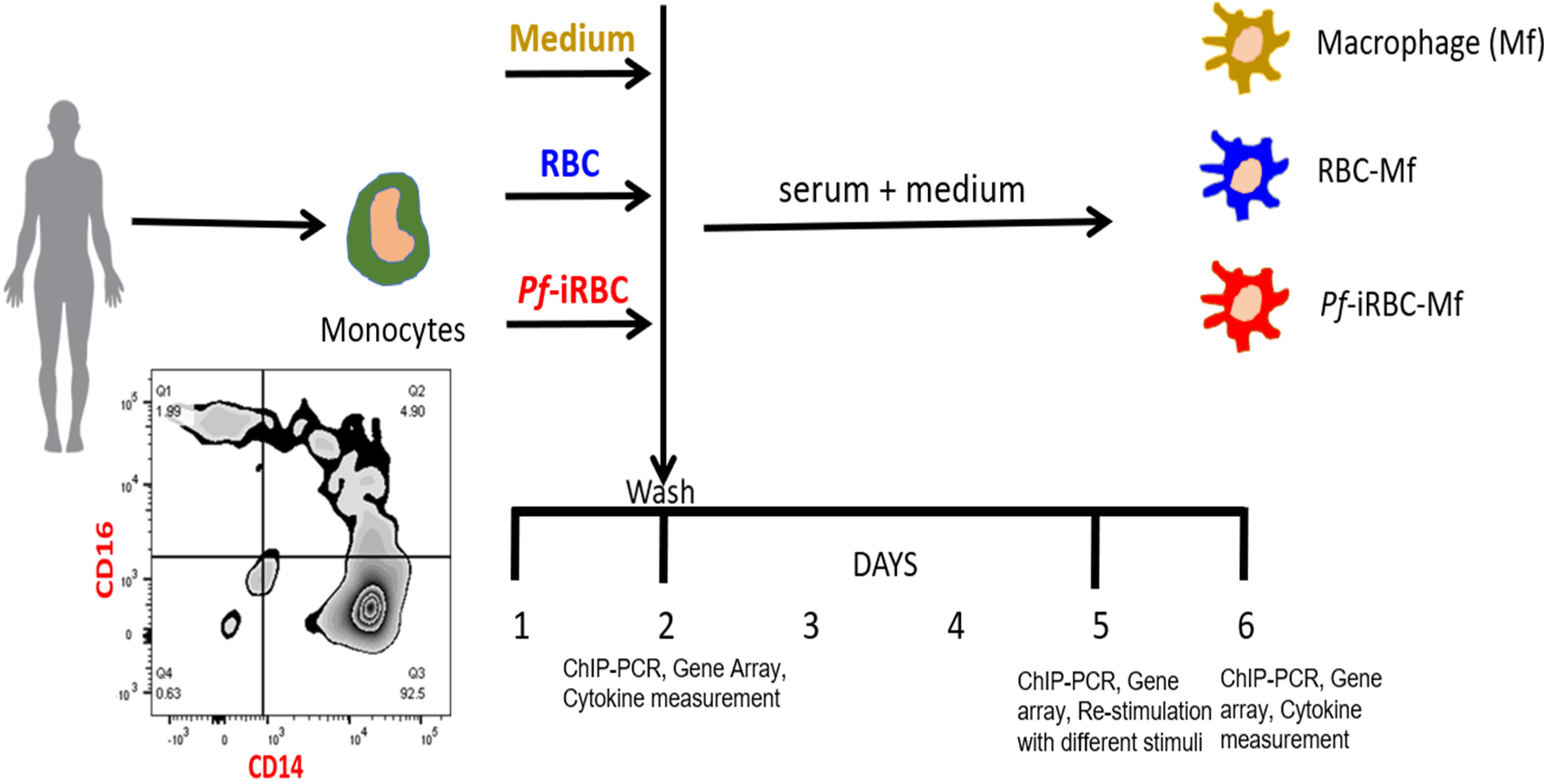
In vitro model of monocyte to macrophage differentiation during exposure to *P*. *falciparum* blood-stage parasites. Elutriated monocytes from healthy U.S. adults were incubated for 24 hours with medium alone, uninfected red blood cells (RBC) or *Pf*-iRBC (monocyte:*Pf*-iRBC ratio 1:15). At 24 hours, supernatants and cells were collected from some replicate wells for cytokine analysis, and the ChIP assay, while monocytes in other replicate wells were washed and incubated for 3 additional days in human serum plus medium to allow monocytes to differentiate into macrophages (Mf). On day 5, the three populations of macrophages (Mf, RBC-Mf and *Pf*-iRBC-Mf) were either harvested for ChIP and cytokine gene expression analysis; or re-stimulated with *Pf*-iRBCs or LPS for 24 hours prior to supernatants and cells being collected for cytokine analysis.

As expected, monocytes of U.S. adults (n=3) co-cultured for 24 hours with *Pf*-iRBCs upregulated the expression of genes encoding several pro-inflammatory cytokines including TNF, IL-6 and IL-1β (Figure 5A), which was confirmed at the protein level in an independent experiment of U.S. adults (n=9) by cytokine analysis of supernatants (Figure 5D-F). On day 5, after monocytes had matured into macrophages, pro-inflammatory cytokine gene expression by *Pf*-iRBC-Mfs decreased relative to the 24-hour timepoint but remained higher than pro-inflammatory cytokine gene expression in the Mf and RBC-Mf controls (Figure 5B), consistent with the removal of *Pf*-iRBC by washing at the 24-hour timepoint. On day 5, Mf, RBC-Mf and *Pf*-iRBC-Mf were co-cultured with *Pf*-iRBC lysate. After 24 hours, gene expression analysis showed upregulation of several pro-inflammatory cytokines in Mf and RBC-Mf relative to the 5-day timepoint that was not apparent in *Pf*-iRBC-Mf (Figure 5C). This pattern was confirmed at the protein level for TNF, IL-6 and IL-1β in an independent experiment of U.S. adults (n=9) by cytokine analysis of supernatants (Figure 5G-I). Plots of ΔCt values (mean ±SE of the 3 subjects in Figure 5A-C) for TNF and IL-6 at the three timepoints (Figure 5J and K) further illustrates how pre-exposure of monocytes to *P*. *falciparum* blunts the subsequent inflammatory response of newly differentiated macrophages upon restimulation with *P*. *falciparum*. We observed a similar dampening of TNF, IL-6 and IL-1β responses to LPS in *Pf*-iRBC-Mf (Figure 5L-N).

**Figure 5.**
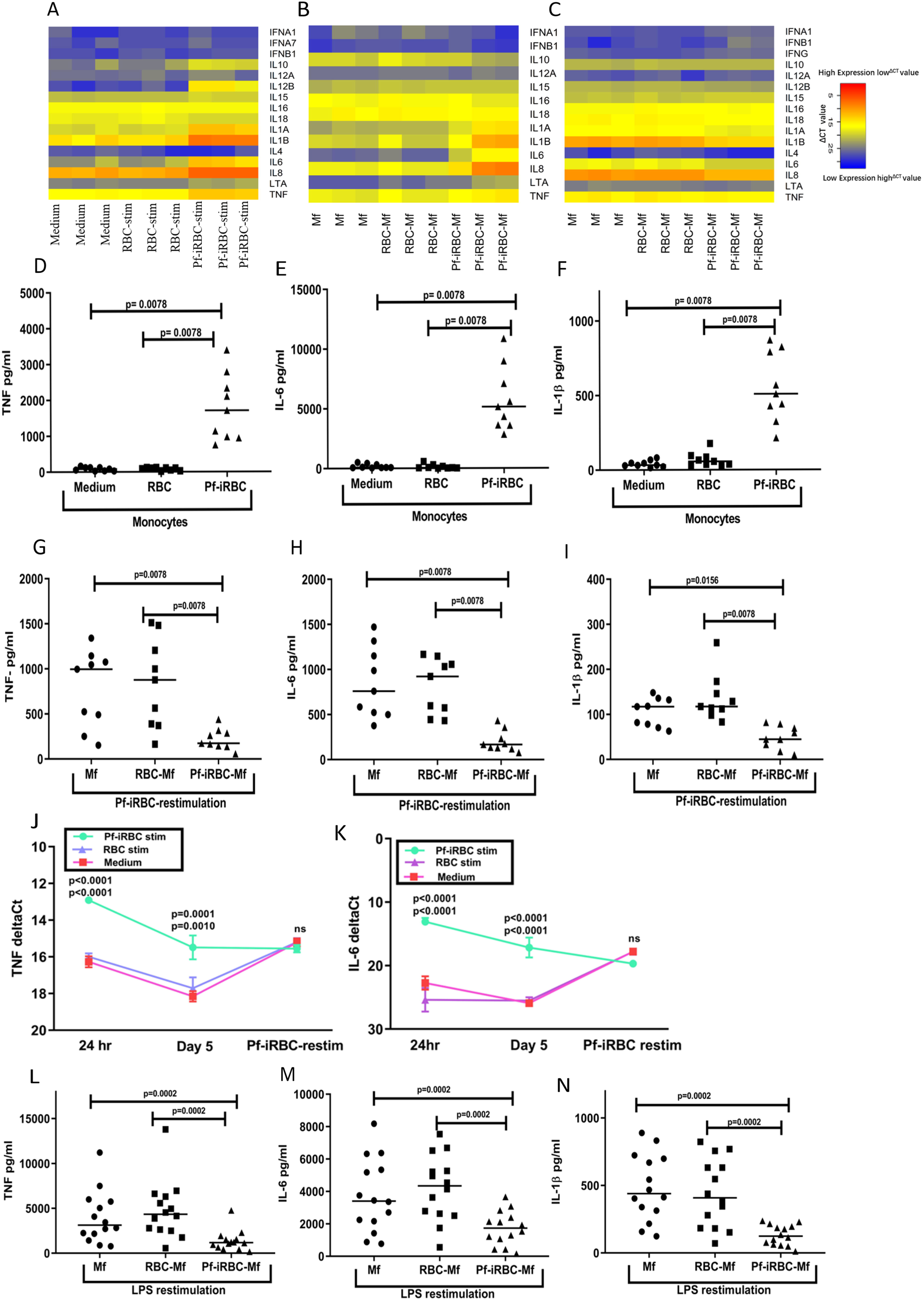
Pre-exposure of monocytes to *P*. *falciparum* dampens subsequent inflammatory responses to *P*. *falciparum* and LPS. Monocytes of U.S. adults were incubated with medium, RBC or *Pf*-iRBC. At 24 hours, cells were analyzed by (**A**) cytokine gene expression Taqman arrays (n=3 subjects), and supernatants were analyzed by a bead-multiplexed assay (n=9 subjects) to quantify (**D**) TNF, (**E**) IL-6 and (**F**) IL-1β. In the same experiment, replicate monocytes were incubated with medium, RBC or *Pf*-iRBC, washed at 24 hours and incubated for 3 additional days in human serum to permit macrophage (Mf) differentiation. On day 5, the three populations of macrophages were analyzed by (**B**) cytokine gene expression arrays (n=3 subjects). Finally, in the same experiment, replicate monocytes were incubated with medium, RBC or *Pf*-iRBC, washed at 24 hours, incubated for 3 additional days in human serum to permit Mf differentiation, and then co-cultured with *Pf*-iRBC or LPS for 24 hours. Cells were harvested for (**C**) cytokine gene expression arrays (n=3 subjects), and supernatants were analyzed (n=9 subjects) to quantify TNF, IL-6 and IL-1β induced by (**G-I**) *Pf*-iRBC or (**L-N**) LPS (n=14 subjects). ΔCt values (mean ±SE) for (**J**) TNF and (**K**) IL-6 at the indicated timepoints for the 3 subjects shown in A-C. (A-C) Heatmaps were generated from ΔCt values, with lower ΔCt values corresponding to higher gene expression. ΔCt values were normalized to 18S rRNA expression. (D-I and L-N) Lines represent median values. Data were analyzed by the Wilcoxon test with Bonferroni adjustment, and significance levels between the groups are indicated by P values. (J and K) Two-way ANOVA was performed followed by Tukey’s multiple comparisons test. Significance level between conditions (Pf-iRBC stim vs. Control Medium and Pf-iRBC stim vs. RBC stim, respectively) are indicated by P values at each timepoint.

### *P*. *falciparum* exposure induces epigenetic changes in monocytes consistent with regulation of inflammation

Epigenetic modifications in monocytes/macrophages underpin the immunological imprinting of tolerance or trained immunity following exposure to LPS or β-glucan, respectively [25]. Here we hypothesized that *P*. *falciparum* induces epigenetic modifications in the regulatory regions of pro-inflammatory genes such that inflammatory responses are dampened upon re-exposure to *P*. *falciparum* (i.e. tolerance). To test this hypothesis, we performed Chromatin Immunoprecipitation (ChIP) on monocytes/macrophages collected at each of the 3 timepoints shown in Figure 4 using an antibody specific for H3K4me3, an epigenetic modification of Histone H3 enriched at active promoters that positively correlates with transcription [26].

After 24-hours, monocytes stimulated with *Pf*-iRBC were enriched for the active H3K4me3 histone mark at the TNF and IL-6 promoter regions relative to monocytes incubated with RBCs or medium alone (Figure 6A, D), consistent with cytokine data at the same timepoint (Figure 5D-F). On day 5, after maturation of monocytes into macrophages (Mf), the active H3K4me3 histone mark remained enriched at the TNF and IL-6 promoter regions of cells that had been stimulated with *Pf*-iRBC (Pf-iRBC-Mf) relative to macrophages initially incubated with RBCs (RBC-Mf) or medium alone (Mf) (Figure 6B, E). After the three cell populations (Mf, RBC-Mf and Pf-iRBC-Mf) were stimulated with Pf-iRBC for 24 hours, the active H3K4me3 histone mark became enriched in the TNF and IL-6 promoter regions of the Mf and RBC-Mf populations, whereas the active H3K4me3 histone mark decreased in the TNF and IL-6 promoter regions in the Pf-iRBC-Mf population (Figure 6C, F). Plots of the H3K4me3 histone mark at the TNF and IL-6 promoter regions at all three timepoints further illustrate how pre-exposure to *P*. *falciparum* diminishes H3K4me3 histone mark enrichment of newly differentiated macrophages upon restimulation with *P*. *falciparum* (Figure 6G and H)—an epigenetic pattern of TNF and IL-6 regulation consistent with the pattern of TNF and IL-6 expression in the same model (Figure 5J and K).

**Figure 6.**
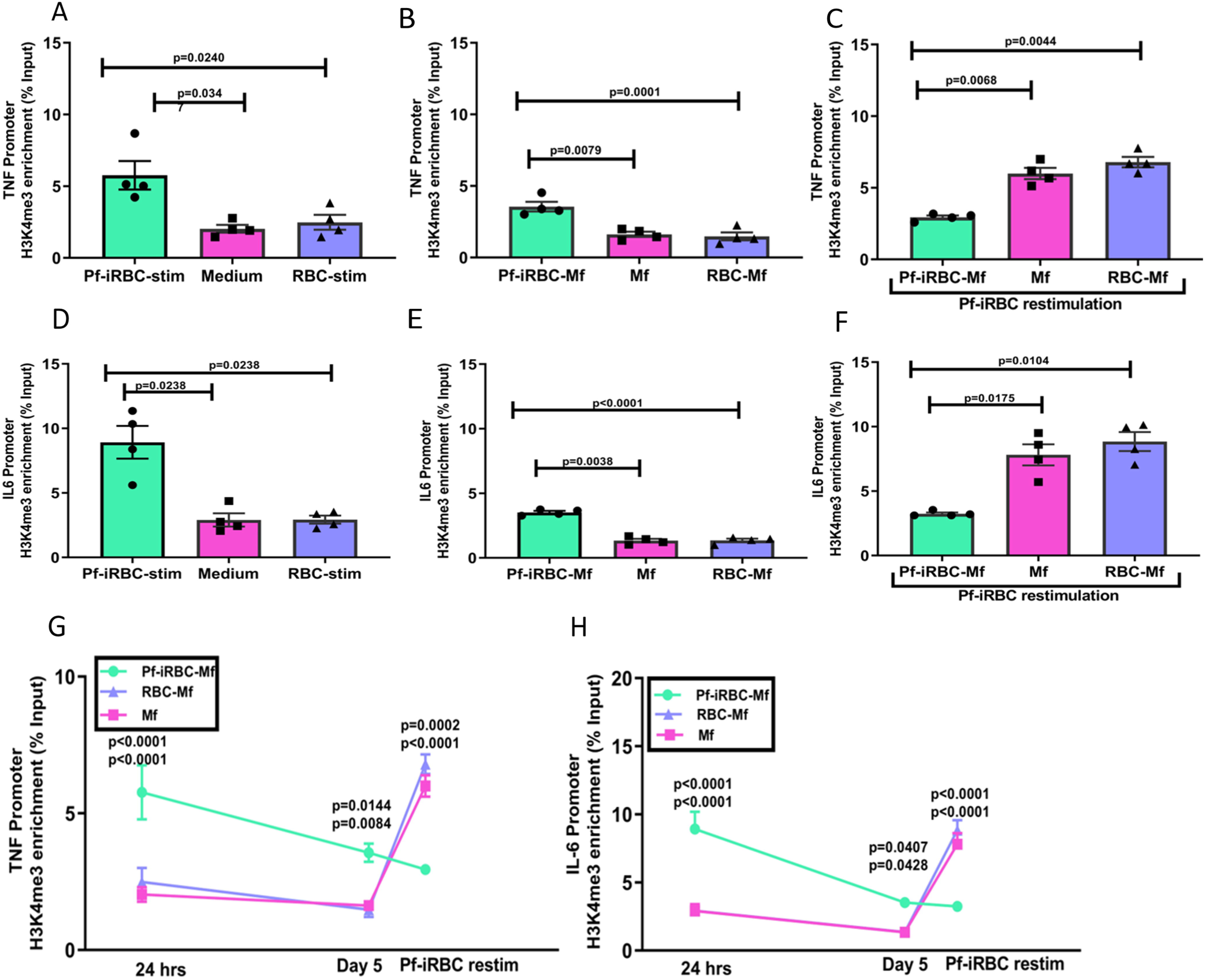
*P*. *falciparum* exposure drives epigenetic reprogramming of monocyte/macrophages toward a tolerant phenotype. Monocytes of U.S. adults (n=4) were incubated with medium, RBC or *Pf*-iRBC for 24 hours and then analyzed by chromatin immunoprecipitation (ChIP) and RT-PCR to quantify H3K4me3 enrichment at (**A**) TNF and (**D**) IL-6 promoter sites. Replicate monocytes were incubated with medium, RBC or *Pf*-iRBC, washed at 24 hours, and incubated for 3 days in human serum to permit Mf differentiation. On day 5, cells were analyzed by ChIP and RT-PCR to quantify H3K4me3 enrichment at (**B**) TNF and (**E**) IL-6 promoter sites. Finally, replicate monocytes were incubated with medium, RBC or *Pf*-iRBC, washed at 24 hours, incubated for 3 days in human serum, co-cultured with *Pf*-iRBC for 24 hours, and analyzed by ChIP and RT-PCR to quantify H3K4me3 enrichment at (**C**) TNF and (**F**) IL-6 promoter sites. Kinetics of H3K4me3 enrichment at (**G**) TNF and (**H**) IL-6 promoter sites across indicated timepoints. Results are shown as means ±SEM fold enrichment of H3K4me3 antibody as percent of input. (A-F) One-way ANOVA with Tukey’s adjustment for multiple comparisons. P values indicate level of significance. (G, H) Two-way ANOVA with Tukey’s adjustment for multiple comparisons. P values indicate level of significance between Pf-iRBC vs. medium and Pf-iRBC vs. RBC, respectively.

## Discussion

In areas of intense malaria transmission the risk of febrile malaria decreases with age as individuals are exposed to repeated *P*. *falciparum* infections over many years [17]. The relatively slow development of clinical immunity to malaria is associated with the gradual acquisition of *P*. *falciparum*-specific adaptive immune responses [27]. Here we sought to understand the impact of *P*. *falciparum* exposure on the phenotype and function of innate immune cells, namely, the monocyte/macrophage lineage—an important source of fever-inducing pro-inflammatory cytokines during blood-stage malaria infection [11].

In the Mali cohort we observed an inverse relationship between age and *Pf*-iRBC-inducible production of the inflammatory cytokines IL-1β, IL-6 and TNF from monocytes; whereas monocytes of Malian infants born during the dry season (negligible P. falciparum transmission) and malaria-naïve U.S. adults produced similarly high levels of inflammatory cytokines, indicating that age alone is not responsible for the functional changes in monocytes observed in the Mali cohort, but rather malaria exposure itself and/or other factors associated with malaria transmission. In addition, monocytes of Malian adults produced higher levels of the anti-inflammatory cytokine IL-10 in response to *Pf*-iRBC stimulation. Consistent with these functional data, monocytes of Malian adults expressed higher levels of the regulatory molecules CD163, CD206, Arginase-1 and TGM2 [28-32].

Importantly, the monocytes analyzed in this study were collected from uninfected, asymptomatic individuals at the end of the 6-month dry season when the immune system is in a relatively non-activated state. Since monocytes survive in circulation for only 1-7 days [15], we postulate that the skewing of circulating monocytes toward a tolerant phenotype with increasing age reflects the ‘reprogramming’ of bone marrow progenitor cells by past malaria exposure and/or other immune perturbations associated with malaria transmission. This hypothesis is consistent with recent studies in mice showing that BCG or β-glucan can epigenetically and metabolically reprogram myeloid progenitors in the bone marrow [33, 34]— a mechanism that may be particularly relevant to malaria as the bone marrow is a major site of growth and sexual development for *Plasmodium* parasites [35]. Alternatlvely, a recent study using the rodent malaria model by Nahrendorf et al. found that epigenetic reprogramming of monocytes in tolerized hosts occurs within the spleen [36].

We previously found in longitudinal analyses of Malian children that *Pf*-iRBC-inducible production of IL-1β and IL-6 by monocytes was lower 7 days after treatment of febrile malaria relative to that induced at the pre-infection baseline before the six-month malaria season [19]. However, the skewing of monocytes toward a tolerant phenotype in children after febrile malaria seems to depend on ongoing *P*. *falciparum* exposure, as children’s monocytes generally appear to return to a ‘non-tolerant’ steady-state baseline after the subsequent 6-month dry season [19]. However, a study in the same cohort found that children who are resistant to febrile malaria begin the malaria season with evidence of tolerized monocytes that upregulate p53, which is associated with attenuation of *Plasmodium*-induced inflammation [37]. Interestingly, a study conducted in a region of Uganda where P. falciparum transmission occurs year-round found that older children had a dampened pro-inflammatory serum cytokine response during acute malaria compared to younger children [38], consistent with the notion that ongoing exposure may be required to maintain a tolerant state in most children.

That innate immune ‘memory’ may be relatively short-lived after a single immune perturbation is consistent with other studies that show the trained immunity phenotype persists for at least 3 months and up to 1 year [13]. Therefore, it seems plausible that with each 6-month malaria season, the homeostatic setpoint of monocytes gradually shifts toward a tolerant phenotype such that adults maintain a tolerant phenotype even through the 6-month dry season. This is consistent with whole blood transcriptome analysis of Malian adults who exhibited a blunted inflammatory response during *P*. *falciparum* infection relative to Dutch adults who were experimentally infected with *P*. *falciparum* for the first time [39].

To more directly assess the impact *P*. *falciparum* blood-stage parasites on monocytes/macrophages and to gain insight into potential molecular mechanisms of *Plasmodium*-induced tolerance, we employed an in vitro system of monocyte to macrophage differentiation and *P*. *falciparum* co-culture [25, 40]. With this model we found at both the mRNA and protein levels that macrophages derived from monocytes previously exposed to *Pf*-iRBCs had an attenuated inflammatory cytokine response upon re-exposure to *Pf*-iRBCs or exposure to the TLR4 agonist LPS. This corresponded to a decrease in the epigenetic marker of active gene transcription, H3K4me3, at inflammatory cytokine gene loci. These findings are consistent with the hypothesis that malaria contributes directly to the reprogramming of monocytes/macrophages toward a tolerant phenotype.

Given the non-specific nature of monocyte tolerance, the findings of this study may have implications for vaccine responsiveness and the clinical course of non-malaria infections in malaria-exposed populations, even when individuals are uninfected and asymptomatic. For example, studies of the PfSPZ malaria vaccine in Mali and Tanzania reported an inverse relationship between cumulative *P*. *falciparum* exposure or transmission intensity and vaccine-specific antibody responses [41, 42]. Given that monocytes can help initiate vaccine-specific adaptive immunity through cytokine production, and to a lesser extent antigen processing and presentation, it is plausible that tolerized monocytes play a role in vaccine hypo-responsiveness in malaria-endemic areas. It is also conceivable that tolerized monocytes/macrophages could modify the clinical manifestations of non-malaria infections, particularly those linked to monocyte/macrophage-associated pathological inflammation. For example, severe acute respiratory syndrome coronavirus 2 (SARS-CoV-2), the causative agent of the coronavirus disease 2019 (COVID-19) pandemic, has been associated with excessive inflammation, including high levels of circulating IL-6 and TNF, that is thought to be a major cause of disease severity and death in patients with COVID-19 [43]. In much of Africa to date, the COVID-19 pandemic has not been as severe as predicted [44]. Although many environmental, genetic, sociocultural and institutional factors could contribute to lower COVID-19 morbidity and mortality in Africa compared to other regions [45], we speculate that COVID-19 severity in Africa may be mitigated by pre-existing differences in the immune system, including tolerized monocytes that produce lower levels of pro-inflammatory cytokines when activated.

In contrast to our findings here, recent studies have reported that malaria induces a state of hyper-responsiveness [40, 46, 47] that is functionally similar to the trained immunity induced by BCG vaccination [13]. For example, Schrum et al. found that initial stimulation with Pf-iRBCs induced human adherent PBMCs to hyper-respond to subsequent stimulation with the TLR2 agonist Pam3CSK4. These findings may differ from the present study for several reasons. In contrast to purified monocytes used in the present study, Schrum et al. performed *in vitro* studies with adherent PBMCs, which may contain cells other than monocyte/macrophages. In addition, our in vitro studies used a higher cell to Pf-iRBC ratio (1:15 vs. 1:0.5) that more closely approximates *in vivo* parasitemia during febrile malaria. Finally, we restimulated monocytes/macrophages with Pf-iRBCs rather than TLR agonists alone.

Taken together, our findings offer mechanistic insight into the long-standing clinical observation that individuals exposed to intense malaria transmission can tolerate malaria parasites in their blood at levels that would predictably produce fever in previously unexposed individuals [16]. In future studies it will be of interest to determine the generalizability of these findings in other malaria-exposed populations and to assess the extent to which variation in transmission intensity influence monocyte phenotype and function. In addition, it will be of interest to track *ex vivo* monocyte epigenetic profiles within individuals over time and with repeated infections to better understand the quality and kinetics of malaria-induced tolerance.

## Materials and Methods

### Study subjects

The field study was conducted in the rural village of Kalifabougou, Mali where intense *P*. *falciparum* transmission occurs from June through December each year. The cohort study has been described in detail elsewhere [17]. Briefly, 695 healthy children and adults aged 3 months to 25 years were enrolled in an ongoing cohort study in May 2011. Exclusion criteria at enrollment included a hemoglobin level <7 g/dL, axillary temperature ≥37.5°C, acute systemic illness, underlying chronic disease, or use of antimalarial or immunosuppressive medications in the past 30 days. The present study focused on children aged 6-months to 8 years and adults. For this study venous blood samples were collected from study subjects at their healthy uninfected baseline before the malaria season. The ethics committee of the Faculty of Medicine, Pharmacy and Dentistry at the University of Sciences, Techniques and Technology of Bamako, and the Institutional Review Board of NIAID NIH approved the study (ClinicalTrials.gov NCT01322581). Written, informed consent was obtained from the parents or guardians of participating children or from adult participants.

### PBMC processing

Blood samples (8 ml) were drawn by venipuncture into sodium citrate-containing cell preparation tubes (BD, Vacutainer CPT Tubes) and transported 20 km to the laboratory where PBMCs were isolated and frozen within three hours according to the manufacturer’s instructions. PBMCs were frozen in human AB serum (Sigma) containing 10% dimethyl sulfoxide (DMSO; Sigma-Aldrich, St. Louis, MO), kept at −80°C for 24 hours, and then stored in liquid nitrogen.

### Isolation of monocytes

PBMCs and elutriated monocytes were obtained from healthy U.S. volunteers by counterflow centrifugal elutriation at the National Institutes of Health (NIH) Blood Bank under Institutional Review Board approved protocols of both the National Institute of Allergy and Infectious Diseases and the Department of Transfusion Medicine. Elutriated monocytes were further purified with a monocyte isolation kit (Stem Cell technologies) to minimize donor-to-donor variability in contaminating cell populations. Monocyte purity was routinely >98% as assessed by flow cytometry. Similarly, monocytes were isolated from PBMCs of Malian donors by a negative selection monocyte isolation kit (Stem Cell Technologies) without depleting CD16 such that monocytes were isolated ‘untouched’. Non-monocyte/macrophage cells were directly depleted with a cocktail of biotin-conjugated antibodies followed by magnetic removal of labeled cells.

### Preparation of *P*. *falciparum*-infected red blood cell lysate for in vitro stimulation

3D7 *P*. *falciparum* parasites were maintained in fresh human ORh^+^ erythrocytes at 3% hematocrit in RPMI 1640 medium (KD Medical) supplemented with 10% heat-inactivated ORh+ human serum (Interstate Blood Bank, Memphis, Tennessee), 7.4% Sodium Bicarbonate (GIBCO, Invitrogen) and 25 mg/ml of gentamycin (GIBCO, invitrogen), at 37°C in the presence of a gas mixture containing 5% O2, 5% CO2 and 90% N2. Parasite cultures were confirmed to be free of mycoplasma and acholeplasma using an ELISA-based Mycoplasma Detection Kit (Roche) which contains polyclonal antibodies specific for *M. arginini, M. hyorhinis, A. laidlawii* and *M. orale. P. falciparum*-infected red blood cells (Pf-iRBCs) were enriched for knobs using Zeptogel (contains gelatin) sedimentation. Pf-iRBCs were enriched at the schizont stage with percoll-sorbitol gradient and centrifugation, washed, and resuspended in complete medium in the absence of human serum or Pf-iRBCs schizonts were isolated in RPMI 1640 medium supplemented with 0.25% Albumax (GIBCO, Invitrogen) and 7.4% Sodium Bicarbonate (GIBCO, Invitrogen) using magnetic columns (LD MACS Separation Columns, Miltenyi Biotec). Control preparations of uninfected red blood cells (RBCs) from the same blood donor were obtained and tested in all experiments. Lysates of Pf-iRBCs and RBCs were obtained by three freeze-thaw cycles using liquid nitrogen and a 37°C water bath.

### In vitro stimulation of PBMCs and monocytes with *P*. *falciparum*-infected red blood cell lysate

Monocytes or PBMCs were cultured with RBCs or Pf-iRBCs. Cells were cultured in complete RPMI (RPMI 1640 plus 10% human AB serum, 1% penicillin/streptomycin), at 37°C in a 5% CO2 atmosphere. PBMCs were stimulated with RBC or Pf-iRBC lystate at a ratio of 3 RBCs or 3 Pf-iRBCs per PBMC, whereas monocytes were stimulated at a ratio of 5-30 RBCs or 5-30 Pf-iRBCs per monocyte. For *in vitro* experiments with elutriated monocytes, cells were first allowed to adhere in monocyte attachment medium (Promocell) for 1.5 hr before culturing as described above.

### Flow cytometry

PBMCs were washed in PBS with 4% heat-inactivated FCS, incubated for 15 min on ice with a live-dead dye in PBS, washed, and then surface stained with lymphocyte lineage dump-APC (CD3, CD19, CD56, CD20), CD14-BUV805, CD206-BV421, CD163-FITC, CD16-BUV395, HLA-DR-APC-R700 and CD86-BV650. For intracellular staining, following surface staining cells were fixed and permeabilized using a Foxp3 staining kit (e-biosciences). Cells were then stained with Arginase1-PE-Cy7 in permeabilization buffer. After washing, cells were resuspended in 4% heat-inactivated FCS containing FACS buffer and data were acquired by a Symphony Flow Cytometer (BD Biosciences). Flow cytometry and t-SNE analyses were performed with FlowJo software (FlowJo10.5.3). For t-SNE analysis, down sampling was done on live monocytes which were devoid of aggregates and dead cells, followed by concatenating the samples according to biological replicates and sample group. Finally, the default t-SNE algorithm was run with 1000 iterations, perplexity 30 and a learning rate of 200.

### Cytokine measurements in supernatants

Supernatants were thawed and immediately analyzed with Bio-plex human cytokine assays (Bio-Rad Laboratories, Inc.) following the manufacturer’s instructions. The following cytokines were measured: IL-1β, IL-6, IL-10 and TNF. Briefly, 50 uL of supernatant was incubated with anti-cytokine antibody-coupled magnetic beads for 30 min at room temperature with shaking at 300 RPM in the dark. Between each step the complexes were washed three times in wash buffer using a vacuum manifold. The beads were then incubated with a biotinylated detector antibody for 30 min before incubation with streptavidin-phycoerythrin for 30 minutes. Finally, the complexes were resuspended in 125 mL of detection buffer and 100 beads were counted with a Luminex 200 device (Bio-Rad Laboratories, Inc.). Final concentrations were calculated from the mean fluorescence intensity and expressed in pg/mL using standard curves with known concentrations of each cytokine.

### RNA isolation, RNA-Seq and cytokine gene expression array

Cells were kept in RNAProtect buffer (Qiagen) at −80°C until RNA was isolated. RNA was isolated using the RNAeasy kit according to the manufacturer’s instructions. The quality and quantity of isolated RNA was determined with the Agilent Bioanalyzer. Only RNA with RIN values greater than 7 were used for analyses. cDNA was prepared from 10 ng of total RNA using the Ovation® RNA-Seq System V2 (Tecan) according to manufacturer’s instructions. This method employs both poly-T and random primers so that both poly-adenylated and non-poly-adenylated RNA is included. The cDNA product was end-repaired using the NEBNext End Repair Module (New England Biolabs). RNA-Seq libraries were prepared using 1 μg of end-repaired cDNA using the TruSeq Stranded RNA Kit (Illumina), however due to the method of amplification the libraries were not stranded. Unique dual-indexed barcode adapters were applied to each library. Libraries were pooled in an equimolar ratio for sequencing. The pooled libraries were sequenced on one lane of an S4 flow cell on a NovaSeq 6000 using version 1 chemistry to achieve a minimum of 49 million 150 base pair reads. The data was processed using RTA version 3.4.4 and BWA-0.7.12. For the RNA-seq analysis, quality control and adapter trimming were performed using FASTQC and cutadapt, respectively (Martin, 2013). Then, the reads were aligned to the hg19 reference genome using STAR aligner (Dobin et al., 2013). Reads were counted using featureCounts (Liao, Smyth, & Shi, 2014). For the identification of differentially expressed genes among the different groups, we used DESeq2 with the design formula “∼ condition” (Love, Huber, & Anders, 2014). Benjamini-Hochberg (BH) correction was performed with an adjusted p-value threshold set to 0.01. For the gene expression array, we used TaqMan™ Array, Human Cytokine Network, fast 96-well plate, and real time PCR was performed from isolated cDNA according to the manufacturer’s instructions. cDNA was isolated from RNA using the Superscript-VILO cDNA synthesis kit followed by Real Time RTPCR using TaqMan™ Fast Advanced Master Mix using the Quant Studio™ 6 instrument.

### Cell culture

For ChIP analysis and Taqman RNA Array analysis, 10×10^6^ elutriated and purified monocytes were plated on 100 mm dishes. Monocytes were pre-incubated with cell culture medium (RPMI), Pf-iRBCs or RBCs for 24 hours in a total volume of 10 mL. After wash-out, cells were cultured in RPMI supplemented with 10% human pooled AB serum containing homeostatic levels of M-CSF that induces macrophage differentiation. Cells were collected at 24 hours and on day 5 were counted prior to chromatin immunoprecipitation. After wash-out, cells were cultured in RPMI supplemented with 10% human pooled AB serum. For cytokines production, 2.5×10^4^ to 5×10^4^ purified or elutriated monocytes were plated in a 96 well flat bottom plate. Monocytes were pre-incubated as above for 24 hours in a total volume of 100-200 µL. After a wash-out, cells were cultured in RPMI supplemented with 10% human pooled serum and supernatants were collected for analysis.

### ChIP analysis

Briefly, cells were fixed in 1% formaldehyde for 10 min and quenched with glycine. Chromatin was sonicated from these cells using a Bioruptor Pico (Diagenode) for four cycles of 10× (30 s ON, 30s OFF) on the HIGH setting. Chromatin precipitation was performed using rabbit anti-human H3K4me3 IgG Ab (Active Motif) as described previously [48]. DNA was then quantified using qPCR with the following primer pairs: IL-6, FW 5⍰-AGCTCTATCTCCCCTCCAGG-3⍰, RV 5⍰-ACACCCCTCCCTCACACAG-3⍰; TNF, FW 5⍰-CAGGCAGGTTCTCTTCCTCT-3⍰, RV 5⍰-GCTTTCAGTGCTCATGGTGT-3⍰[40]. For all ChIP experiments, qPCR values were normalized as percent recovery of the input DNA.

### Statistical analysis

Most continuous data were compared using the unpaired Mann-Whitney test or paired Wilcoxon sign rank test, as appropriate. Bonferroni adjustments were applied to correct for multiple comparisons where needed. One-way or two-way ANOVA with Tukey post hoc tests were used to compare continuous variables in situations where data was assumed to be normal or where the two-way experimental design precluded a nonparametric test. All statistical tests are specified in the figure legends. Statistical significances were defined using 2-tailed p-values or adjusted p-values of 0.05 or less. Most statistical tests were computed using GraphPad Prism version 8 (http://www.graphpad.com/scientific-software/prism/). Some heatmaps, principal components analysis (PCA) plots and t-SNE plots were produced using R 3.6.1 or FlowJo (version 10.5)

### Geo Accession ID

GSE151116

## Supporting information

Supplemental Figue-1

## Acknowledgments

We thank the residents of Kalifabougou, Mali, for participating in this study. We also thank Dr. Alice Young and other members of the National Human Genome Research Institute for the RNA sequencing. This work was supported by the Division of Intramural Research of the National Institute of Allergy and Infectious Diseases, National Institutes of Health.

## Figure Legends

**Supplementary Table:** Table of differentially expressed genes of Pf-iRBC-stimulated PBMCs of Malian adults versus children.

This table shows statistically significant (padj<0.01) differentially expressed genes in monocytes from Malian adults stimulated with Pf-iRBC relative to Pf-iRBC stimulated monocytes from Malian children. Adjusted P values were calculated with the Benjamini-Hochberg method.

**Supplementary Figure 1. Titration of monocyte:*Pf*-iRBC ratio for vitro model of monocyte to macrophage differentiation**. Elutriated monocytes from healthy U.S. adults (n=3) were co-cultured with increasing concentrations of *Pf*-iRBCs. After 24 hours, IL-1β (**A**), TNF (**B**) and IL-6 (**C**) were measured in supernatants, and (**D**) cell viability was determined by trypan blue dye exclusion and expressed as percent viability.

**Supplementary Figure 2. Cell viability at day 5 in the vitro model of monocyte to macrophage differentiation**. Elutriated monocytes from healthy U.S. adults (n=9) were incubated for 24 hours with medium alone, uninfected red blood cells (RBC) or *Pf*-iRBC (monocyte:*Pf*-iRBC ratio 1:15). At 24 hours, cells were washed and incubated for 3 additional days in human serum plus medium to allow monocytes to differentiate into macrophages (Mf). To quantify cell viability on day five, 10% v/v alamarBlue HS was added to the culture medium of the three populations of macrophages (Mf, RBC-Mf and *Pf*-iRBC-Mf) for 5 hours and fluorescence intensity (FI) was measured according to the manufacturer’s instructions. FI was normalized to the fluorescence signal in media without cells. Data were analyzed by the Wilcoxon test with Bonferroni adjustment, and levels of significance between the groups are indicated.

