## Supplementary figures and images for "*Plasmodium falciparum* malaria drives epigenetic reprogramming of human monocytes toward a regulatory phenotype"

### Supplemental Figue-1

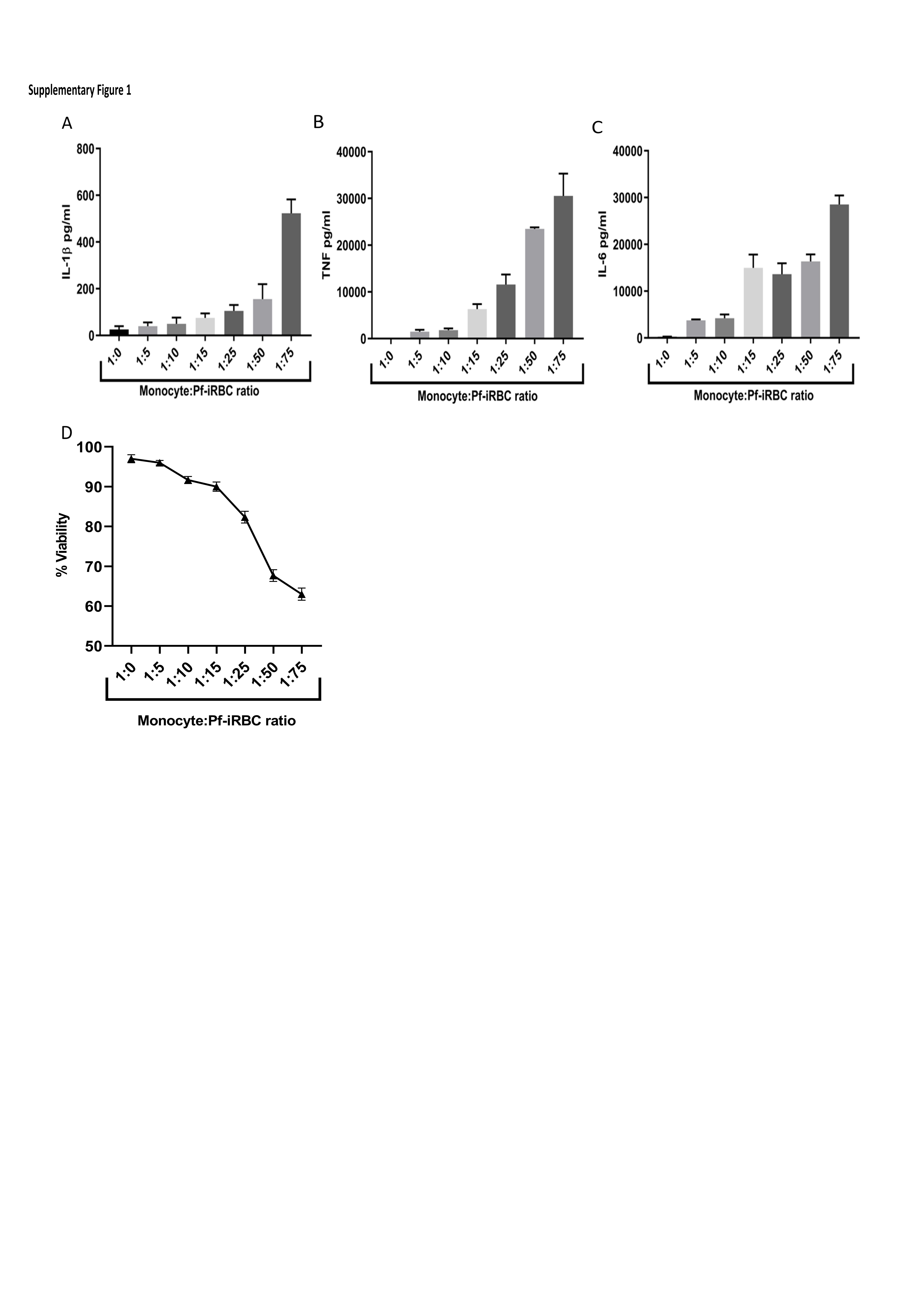

### Supplemental Figure-2

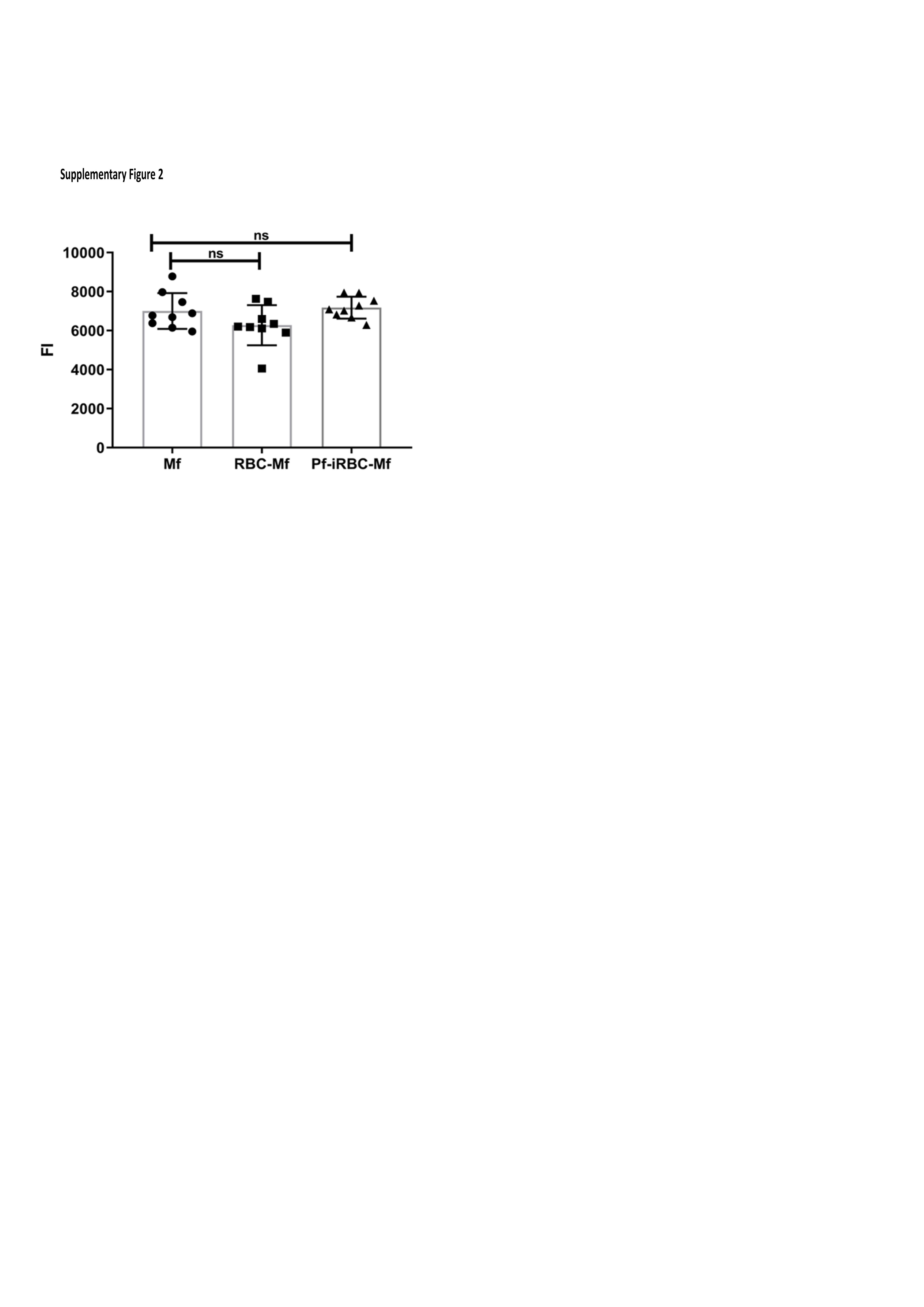
